# Representation of speech in noise in the aging midbrain and cortex: aging may dominate over hearing-loss

**DOI:** 10.1101/212159

**Authors:** Alessandro Presacco, Jonathan Z. Simon, Samira Anderson

## Abstract

**Objective:** To understand the effect of peripheral hearing loss on the representation of speech in noise in the aging midbrain and cortex.

**Methods:** Subjects comprised 17 normal-hearing younger adults, 15 normal-hearing older adults and 14 hearing-impaired older adults. The midbrain response, measured with Frequency-Following Responses (FFRs), and the cortical response, measured with magnetoencephalography (MEG) responses, were recorded from subjects listening to speech in quiet and noise at varying signal to noise ratios (SNRs).

**Results:** Both groups of older listeners showed both weaker midbrain response amplitudes and overrepresentation of cortical responses compared to younger listeners. However, significant differences between the older groups were found in both midbrain-cortex relationships and in cortical processing durations, suggesting that hearing loss may alter reciprocal connections between lower and higher levels of the auditory pathway.

**Conclusions:** The paucity of differences in midbrain or cortical responses between the two older groups suggest that age-related temporal processing deficits may contribute to older adults’ communication difficulties beyond what might be predicted from peripheral hearing loss alone.

**Significance:** Clinical devices, such as hearing aids, should not ignore age-related temporal processing deficits in the design of algorithms to maximize user benefit.

**HIGHLIGHTS:**

- Mild sensorineural hearing loss does not appear to significantly exacerbate already appreciable age-related deficits in midbrain speech-in-noise encoding.
- Mild sensorineural hearing loss also does not appear to significantly exacerbate already appreciable age-related deficits in most measures of cortical speech-in-noise encoding.
- Central processing deficits caused by peripheral hearing loss in older adults are seen only in more subtle measures, including altered relationships between midbrain and cortex.

## 1. Introduction

Speech understanding significantly degrades with aging, particularly in noisy environments. Behavioral studies have suggested that deficient auditory temporal processing is a key factor in explaining age-related difficulties in understanding speech in noise (Pichora-Fuller et al., 1991, Fitzgibbons et al., 1996, Frisina et al., 1997, Schneider et al., 1999, Fitzgibbons et al., 2001, Gordon-Salant et al., 2006, He et al., 2008). Several electrophysiological studies have indeed suggested that these communication problems are linked to temporal processing deficits arising from subcortical (Caspary et al., 1995, Walton et al., 1998, Caspary et al., 2005, Caspary et al., 2006, Parthasarathy et al., 2010, Parthasarathy et al., 2011, Anderson et al., 2012, Presacco et al., 2015, Ananthakrishnan et al., 2016, Presacco et al., 2016a, b) and cortical regions (Soros et al., 2009, de Villers-Sidani et al., 2010, Hughes et al., 2010, Juarez-Salinas et al., 2010, Ross et al., 2010, Lister et al., 2011, Alain et al., 2014, Getzmann et al., 2015, Getzmann et al., 2016, Overton et al., 2016, Presacco et al., 2016a, b, Maamor et al., 2017). Peripheral deficits may also play an important role in altering the final representation of the auditory object, as a progressive loss of cochlear synapses and nerve fibers have been observed in aging animal models (Schmiedt et al., 1996, Sergeyenko et al., 2013). Additionally, recent results have proposed the possibility that decreased cortical connectivity may contribute to speech understanding difficulties (Peelle et al., 2010). This loss of connectivity may cause reduced coherence among different areas of the brain involved in speech comprehension, such as inferior and middle frontal gyrus, along with an unusual activation of larger areas of the brain, including the cyngular-opercular network and the dorsal prefrontal cortex (Wong et al., 2010, Vaden et al., 2015). Additionally, speech understanding in adverse conditions improves when there is strong functional connectivity between the left angular gyrus and medial and left lateral prefrontal cortices (Obleser et al., 2007).

While the findings of these aging studies are quite compelling, they sidestep issues arising from concomitant peripheral hearing loss that often accompanies aging and compromises speech understanding (Humes et al., 1990, Humes et al., 1991). A number of studies have shown that decreased audibility affects auditory temporal processing (Humes et al., 1990, Humes et al., 1991, Anderson et al., 2013, Henry et al., 2014, Petersen et al., 2015, Ananthakrishnan et al., 2016, Maamor et al., 2017, Millman et al., 2017) and leads to reorganization of cortical activity and changes in cortical resource allocation (Peelle et al., 2011, Campbell et al., 2013, Du et al., 2016, Vaden et al., 2016). This degraded auditory temporal processing associated with age-related peripheral hearing loss may contribute to the observation that the use of hearing aids often fails to improve speech understanding in noise. Hearing aid programming focuses on increasing audibility within the auditory dynamic range of the listener, but greater audibility may not restore temporal precision degraded by aging.

Results from our previous studies (Presacco et al., 2016a, b) conducted on normal-hearing older adults showed a substantial degradation (e.g. significantly lower amplitude) of the midbrain response and an exaggeration (or overrepresentation) of the cortical response, both in quiet and noise, with respect to normal hearing younger adults. Overall, our results suggest that temporal processing deficits in the central auditory system contribute to speech-in-noise problems experienced by older adults. Peripheral hearing loss, however, could not be ruled out as a contributing factor. Despite having clinically normal audiometric thresholds, the older adults in our previous study had significantly worse hearing thresholds than the younger adults at most of the frequencies tested. To investigate the role of peripheral hearing loss, we expanded the scope of our previous studies by testing hearing impaired older (OHI) adults.

Our main hypothesis is that the presence of hearing loss exacerbates speech understanding difficulties but does not significantly alter the age-related temporal processing deficits that we observed in normal hearing (ONH) older adults. Specifically, we hypothesize that the Frequency-Following Response (FFR) recorded from the midbrain will be equally degraded (e.g. reduced amplitude with respect to normal hearing younger adults) in ONH and OHI adults. Similarly, in the cortex we expect the reconstruction fidelity, or the ability of the brain to track the speech envelope, to be equally overrepresented in both ONH and OHI adults. Finally, based on the results from our previous study that suggested that the overrepresentation of the cortical response is a negative rather than positive effect of aging (Presacco et al., 2016a), we expect to find a negative correlation between cognitive performance and cortical responses in both groups of older adults.

## 2. Materials and methods

### 2.1 Participants

Participants comprised the same 17 younger normal-hearing (YNH) adults (18 – 27 years, mean ± sd 22.23 ± 2.27, 3 male) and 15 older normal-hearing (ONH) adults (61 – 73 years old, mean ± sd 65.06 ± 3.30, 5 males) that we used for our previous studies (Presacco et al., 2016a, b) and an additional 14 older hearing-impaired (OHI) adults (62 – 86 years old, mean ± sd 71.28 ± 6.26, 9 males) recruited from the Maryland, Washington D.C. and Virginia areas. The OHI group was significantly older than the ONH group (*p* = 0.002). All procedures were reviewed and approved by the Institutional Review Board (IRB) of the University of Maryland. Participants gave informed consent and were paid for their time. YNH and ONH adults had clinically normal audiometric thresholds defined as follows: (1) air conduction thresholds ≤ 25 dB HL from 125 to 4000 Hz bilaterally; and (2) no interaural asymmetry (> 15 dB HL difference at no more than two adjacent frequencies). OHI adults had sensorineural hearing loss with average pure-tone thresholds from 500 – 4000 Hz ≥ 26 dB HL with no thresholds in this frequency range > 90 dB HL. No air-bone gaps greater than 10 dB were noted at any frequency. Figure 1 shows average audiograms for the three groups. All participants from the three groups had normal IQ scores [≥ 85 on the Wechsler Abbreviated Scale of Intelligence (Zhu et al., 1999)] and were not significantly different on IQ (F_[2,43]_ = 0.429, *p* = 0.654). ONH and OHI adults were also not significantly different in sex (Fisher’s exact, *p* = 0.143). Both groups of older adults were screened for dementia on the Montreal Cognitive Assessment (MOCA) (Nasreddine et al., 2005). The mean ± SD of the dementia screening was 26.9 ± 2.7 for ONH and 26.7 ± 2 for OHI adults and no significant differences were found between the two older listener groups (F_[1,27]_ = 0.027, *p*= 0.871). All our participants included in this study scored at our screening criterion of 22 or above. The Edinburgh Handedness Inventory was also administrated to our participants to assess their right or left hand dominance. All the participants, but two YNH and one ONH, were right-handed. Because of the established effects of musicianship on subcortical auditory processing (Bidelman et al., 2010, Parbery-Clark et al., 2012), professional musicians were excluded. All participants participated in both the EEG and MEG study and spoke English as their first language. EEG and MEG data for each participant were collected in two separate sessions.

**Fig 1.**
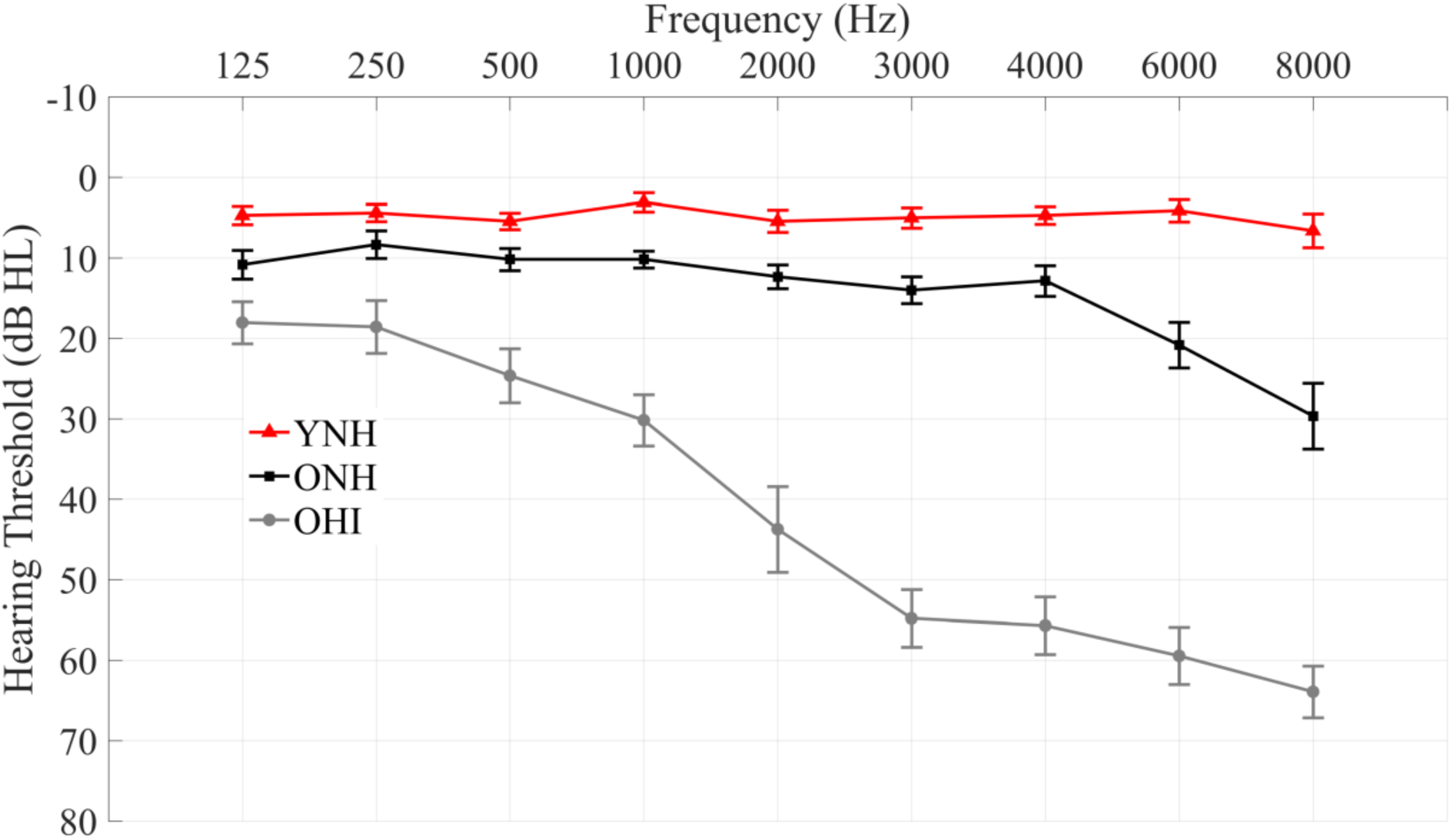
Audiogram (mean ± 1 SE) of the grand averages (right and left) of YNH (red), ONH (black) and OHI (blue) adults. YNH and ONH have pure-tone averages ≤ 25 dB HL from 125 to 4000 Hz, while OHI have an average hearing loss across 500 – 4000 Hz of 26 dB HL or worse.

### 2.2 Speech Intelligibility

The Quick Speech-in-Noise test (QuickSIN) (Killion et al., 2004) was used to quantify the ability to understand speech presented in noise composed of four-talker babble. Speech was presented at a sound level of 70 dB HL to all of our participants, except for two of the participants with hearing loss who required 80 dB HL to ensure proper audibility.

### 2.3 EEG: Stimuli and recording

A 170-ms /da/ (Anderson et al., 2012) was synthesized at a 20 kHz sampling rate with a Klatt-based synthesizer (Klatt, 1980). The stimulus was presented at 75 peak dB SPL diotically with alternating polarities at a rate of 4 Hz through electromagnetically shielded insert earphones (ER-1; Etymotic Research) via Xonar Essence One (ASUS) using Presentation (Neurobehavioral Systems, Inc.). FFRs were recorded in quiet and in 4 noise levels at +3, 0, -3, and -6 signal-to-noise ratios (SNRs) as described in Presacco et al. (2016b). The EEG data were recorded at a sampling frequency of 16384 Hz using the Biosemi ActiABR-200 acquisition system (BioSemi B.V.) using the same montage and filter specifications as described in Presacco et al. (2016b). During the recording session (~2 hr), participants sat in a recliner and watched a silent, captioned movie of their choice to facilitate a relaxed yet wakeful state. Two thousand sweeps that were free of artifacts of physiological origin were recorded for each of the nine total noise levels from each participant.

### 2.4 EEG: data analysis

Data recorded with Biosemi were analyzed in MATLAB (MathWorks, version R2011b) after being converted into MATLAB format with the function pop_biosig from EEGLab (Delorme et al., 2004). The offline analysis of the data was the same as the one described in Presacco et al. (2016b). Sweeps of both polarities were added to minimize the influence of cochlear microphonic and stimulus artifact on the response and to maximize the envelope response (Gorga et al., 1985, Aiken et al., 2008, Campbell et al., 2012).

### 2.5 MEG recording

The same participants in the EEG study also participated in the MEG experiment. The task and stimuli were the same as the ones described in Presacco et al. (2016a). The sound pressure level (SPL) delivered to normal hearing participants was approximately 70 dB SPL when presented with a solo speaker to the participants’ ears with 50 Ω sound tubing (E-A-RTONE 3A; Etymotic Research), attached to E-A-RLINK foam plugs inserted into the ear canal. The level was adjusted upward for hearing impaired participants, if necessary, to ensure audibility. This adjustment was done with noise-free speech and the final sound level was determined by gradually increasing the loudness of the speech until the participant reported the stimulus to be sufficiently audible to perform the task. The entire acoustic delivery system was equalized to give an approximately flat transfer function from 40 to 3000 Hz, thereby encompassing the range of the presented stimuli. Neuromagnetic signals were recorded using a 157-sensor whole head MEG system (Kanazawa Institute of Technology, Kanazawa, Japan) in a magnetically shielded room as described in Ding et al. (2012). A set of 50 ms (10 ms cosine ramped), 500 Hz tones was also presented to each participant at the end of each study. Subjects were asked to count the number of occurrences of the tone (100 sweeps).

### 2.6 MEG: Data analysis

The same data analysis described by Presacco et al. (2016b) was applied to the data. The noise floor was calculated by using the neural response recorded from each noise level tested to reconstruct the speech envelope of a different stimulus than was used during that response. The different stimulus used was a 1-minute speech segment extracted from a different story than that used here, to allow the noise floor to incorporate contributions from potential overfitting.

### 2.7 Cognitive test

The Flanker Inhibitory Control and Attention Test of the National Institutes of Health Cognition Toolbox was used to measure executive function (ability to inhibit visual attention to irrelevant tasks) and attention. Participants were shown a series of arrows and were asked to determine as quickly as possible the direction of the middle arrow by pressing the space bar. The unadjusted scale score was used to compare age-related differences. One OHI subject was removed from all the correlation analyses that involved the cognitive score, as her age was 86 years and no normative values for the cognitive test are available for individuals older than 85.

### 2.8 Statistical analyses

All statistical analyses were conducted in SPSS version 21.0 (SPSS). Fisher’s z transformation was applied to all the correlation values calculated for the midbrain and MEG analyses before running statistical analyses. Repeated-measures ANOVAs were performed to investigate noise level (noise, 4 levels: + 3 dB, 0 dB, -3 dB and -6 dB signal-to-noise ratio) × group interactions in both MEG and FFR data. Split plot ANOVAs were used to test for age group × noise interactions for the RMS values of the FFR response in the time domain and for correlation values calculated for the MEG data. The Greenhouse-Geisser test was used when the Mauchly’s sphericity test was violated. One-way analyses of variance (ANOVAs) were used to analyze between-group differences in FFR amplitude RMS values, for FFR correlation values and for MEG correlation values. The non-parametric Mann-Whitney U and Kruskal-Wallis H tests were used in place of one-way ANOVA when Levene’s test of Equality of Variances was violated. Two-tailed Spearman’s rank correlation (ρ) was used to evaluate the relationships among cognitive score and midbrain and cortical parameters. The false discovery rate (FDR) procedure (Benjamini et al., 1995) was applied to control for multiple comparisons where appropriate.

## 3. RESULTS

### 3.1 Speech Intelligibility

The Kruskal-Wallis test showed significant differences in QuickSIN results among the 3 groups (χ^2^ = 27.566, *p* < 0.001). Post-hoc Mann-Whitney *U* tests showed that YNH (mean ± SD = -0.57 ± 1.13 dB SNR loss) performed significantly better than ONH (mean ± SD = 0.8 ± 1.25 dB SNR loss) (*p* = 0.002) and OHI (mean ± SD = 4.42 ± 3.23 dB SNR loss) (*p* < 0.001). OHI also performed significantly worse than ONH (*p* < 0.001).

### 3.2 Midbrain (EEG): Amplitude analysis

Figure 2 shows the grand average of FFRs to the stimulus envelope of YNH, ONH and OHI in quiet and the most severe noise condition (-6 dB SNR). The ability of midbrain neurons to synchronize in response to the stimulus was assessed by measuring the strength of the FFR via its RMS value. Overall results show a stronger response in younger adults in both the transition (18 – 68 ms) and steady-state (68 – 170 ms) regions. Older adults’ responses show evidence of degradation even in the quiet condition and are not much more in the noise conditions. The presence or absence of hearing loss in older adults does not significantly affect the strength of the response. Details follow below. Figure 3 displays the FFR RMS values for YNH, ONH and OHI for every SNR level tested.

**Fig 2.**
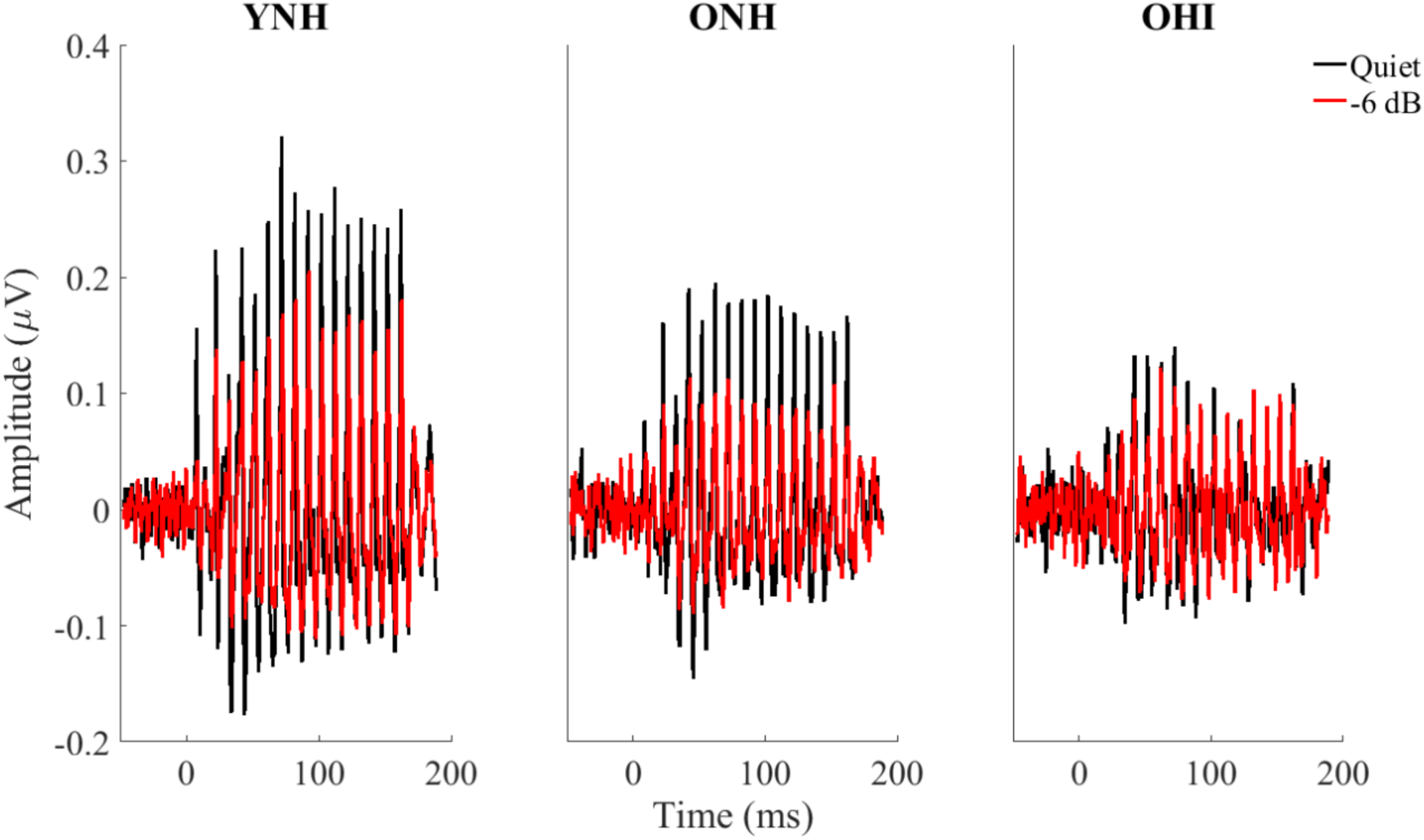
Grand averages for YNH (n = 17, left column), for ONH (n = 15, middle column) and for OHI (n = 14, right column) of FFRs to the stimulus envelope recorded in quiet (black) vs -6 dB noise (red). Overall results show a stronger response in younger adults in both the transition (18 – 68 ms) and steady-state (68 – 170 ms) regions. The presence or absence of hearing loss in older adults does not significantly affect the strength of the response.

**Fig 3.**
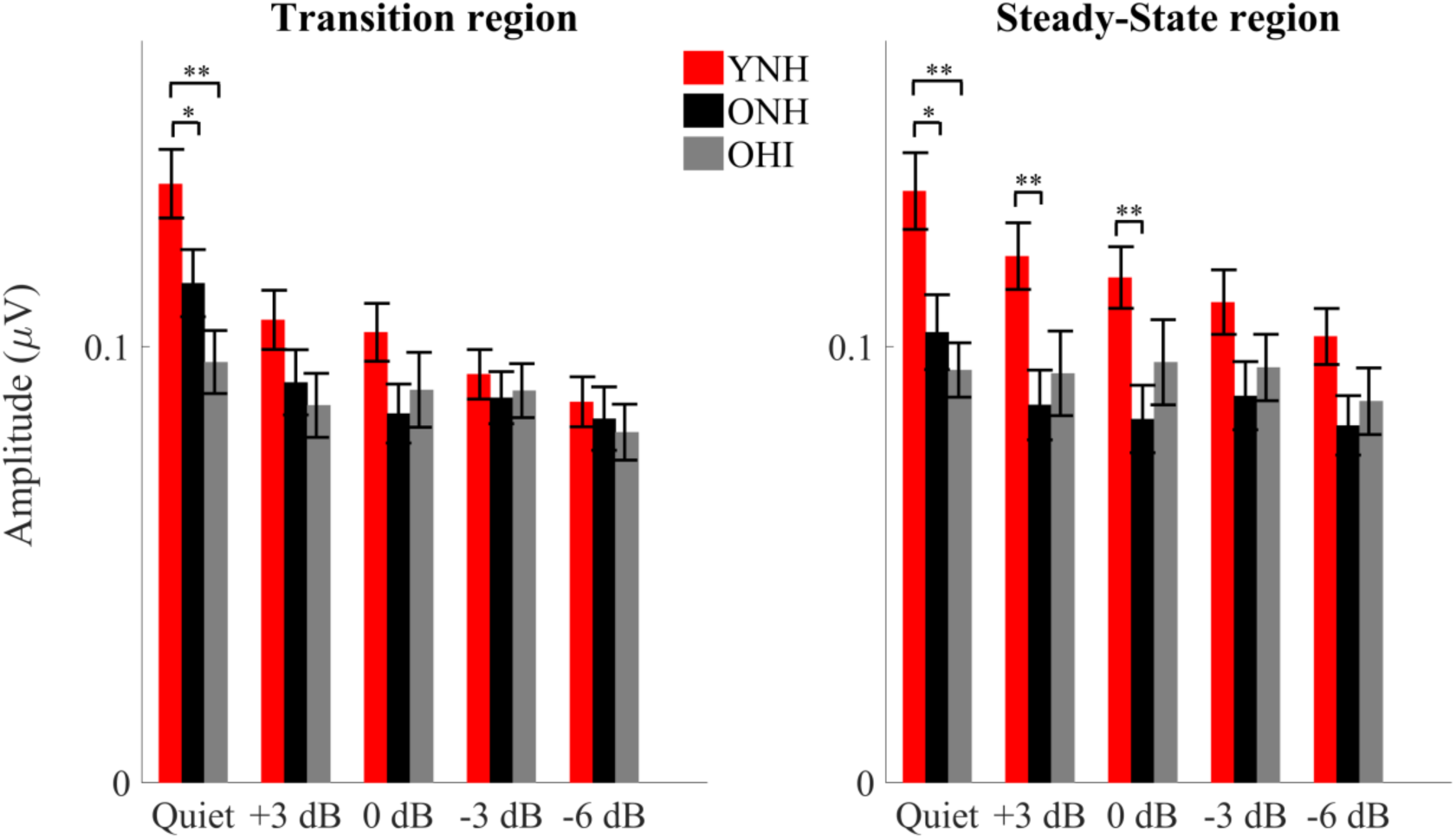
RMS values ± 1 SE for ONH (black) and OHI (Blue) adults in the transition (*left*) and steady-state (*right*) regions for all of the noise levels tested. Higher amplitudes were noted in the quiet condition in the YNH vs. either ONH or OHI. A steeper amplitude decline from quiet to noise conditions was noted in YNH compared to ONH or OHI groups. \**P* < 0.05, \*\**P* < 0.01

*Transition region.* Quiet: A one-way ANOVA showed significant differences across the three groups in the quiet condition (F_[2,43]_ = 7.238, *p* = 0.002). Post-hoc t-tests showed larger amplitudes in YNH than in ONH (t_[30]_ = 2.063, *p* = 0.048) and larger amplitudes in YNH than in OHI (t_[29]_ = 3.758, *p* = 0.001), but not between ONH and OHI (t_[27]_ = 1.707, *p* = 0.099). Noise: A Multivariate ANOVA showed no significant differences among the three groups across the four noise conditions (F_[2,43]_ = 1.326, *p* = 0.242). Repeated measures ANOVA including the quiet and four noise conditions showed a significant condition × age interaction (F_[2,43]_ = 5.486, *p* < 0.001), driven by a steeper amplitude decline from quiet to noise conditions in the YNH than in the ONH or OHI groups, as seen in Figs. 2 and 3. When comparing the noise levels only, repeated measures ANOVA showed no noise level × age interaction (F_[2,43]_ = 2.040, *p* = 0.079).

*Steady-state region.* Quiet: A one-way ANOVA showed significant differences among the three groups in quiet (F_[2,43]_ = 7.351, *p* = 0.002). Post-hoc t-tests showed larger amplitudes in YNH than in ONH (t_[30]_ = 2.622, *p* = 0.014) and larger amplitudes in YNH than in OHI (t_[29]_ = 3.662, *p*= 0.001), but not between ONH and OHI (t_[27]_ = 0.809, *p* = 0.426). Noise: A Multivariate ANOVA showed no significant differences among the three groups across the four noise conditions (F_[2,43]_ = 1.635, *p* = 0.128). Similar to the transition region, Repeated measures ANOVA including the quiet and four noise conditions showed a significant condition × age interaction (F_[2,43]_ = 3.223, *p* = 0.003), driven by a steeper amplitude decline from quiet to noise conditions in the YNH than in the ONH or OHI groups (Figs. 2 and 3). When comparing the noise levels only, repeated measures ANOVA showed no noise level × age interaction (F_[2,43]_ = 1.906, *p* = 0.100).

### 3.3 Midbrain (EEG): Quiet-to-Noise correlation analysis

In order to analyze the robustness of the response profile in noise (the degradative effect of noise), we linearly correlated (Pearson correlation) the average response (Figure 4) obtained in quiet with the ones obtained in noise, for both the transition and steady-state regions for each participant.

**Fig 4.**
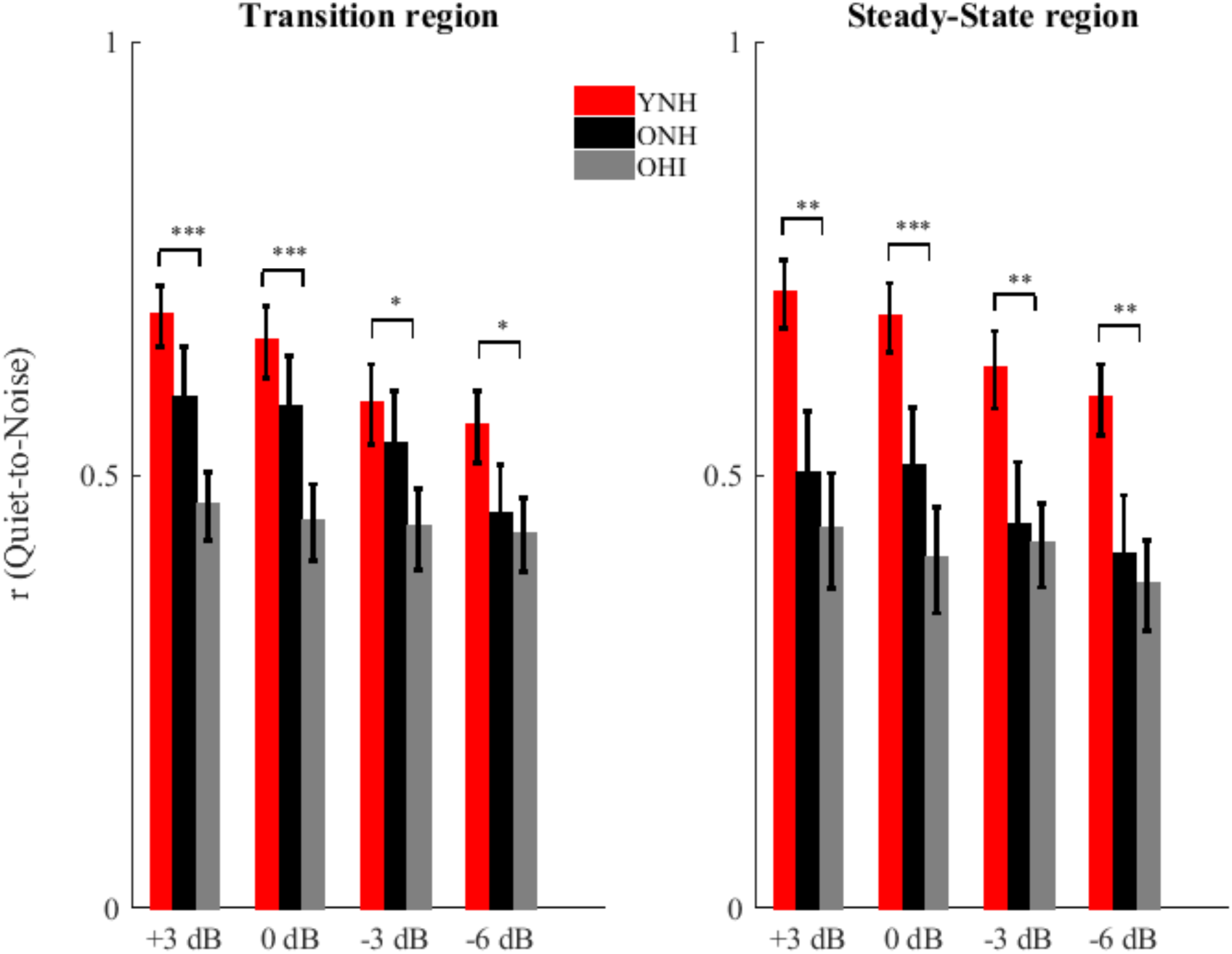
Pearson correlation coefficient ± 1 SE of the quiet-to-noise correlation for YNH (red), ONH (black) and OHI (gray) in the transition (left) and steady-state (right) regions for all of the noise levels tested. Significant differences were noted between YNH and OHI but not between YNH and ONH or ONH and OHI.

*Transition region:* There was a main effect of group across SNRs (F_[2,43]_ = 3.924, *p* = 0.027). Post-hoc one-way ANOVA showed significantly stronger quiet-to-noise correlations in YNH than in OHI (F_[1,30]_ = 3.661, *p* = 0.025), but no differences between YNH and ONH (F_[1,31]_ = 1.344, *p* = 0.280) or between ONH and OHI (F_[1,28]_ = 1.635, *p* = 0.128). There was a significant condition × group interaction (F_[2,43]_ = 3.230, *p* = 0.005) that was driven by a steeper decline in correlation values with decreasing SNR in YNH vs. OHI (F_[1,30]_ = 3.661, *p* = 0.025) that was not seen between YNH and ONH (F_[1,32]_ = 1.344, *p* = 0.280) or between ONH and OHI (F_[1,28]_ = 2.535, *p* = 0.080). Repeated-measures ANOVA showed significant differences across noise conditions in YNH (F_[1,16]_ = 10.101, *p* = 0.001) and ONH (F_[1,14]_ = 8.908, *p* = 0.002), but not in OHI (F_[1,13]_ = 0.366, *p* = 0.779).

*Steady-state region:* There was a main effect of group across SNRs (F_[2,43]_ = 5.207, *p* = 0.009). Post-hoc one-way ANOVA showed significant larger correlations in YNH than in OHI (F_[1,30]_ = 13.823, *p* = 0.001), but not between YNH and ONH (F_[1,31]_ = 3.999, *p* = 0.055) or between ONH and OHI (F_[1,28]_ = 0.853, *p* = 0.364). There was no significant condition × group interaction (F_[2,43]_ = 2.198, *p* = 0.059).

### 3.4 Midbrain (EEG): Stimulus-to-response correlation

The correlation between the clean stimulus and neural response was calculated in order to quantify the ability of the brain to follow the auditory input. No differences were observed among the groups in the quiet condition, but in noise, the YNH group had significantly higher stimulus-to-response correlations than the ONH or OHI groups.

*Quiet:* One-way ANOVA showed no significant differences among the three groups in quiet (F_[2,43]_ = 2.840, *p* = 0.069).

*Noise:* Repeated measures ANOVA showed significant differences among groups across the noise conditions (F_[2,43]_ = 5.288, *p* = 0.009). Follow-up ANOVA showed higher correlation values in YNH compared to either ONH (F_[1,31]_ = 8.651, *p* = 0.006) or OHI (F_[1,30]_ = 6.901, *p* = 0.014) but not between ONH and OHI (F_[1,28]_ = 0.041, *p* = 0.841). The condition × group interaction was not significant (F_[2,43]_ = 0.309, *p* = 0.931).

### 3.5 Cortex (MEG): M100 amplitude and reconstruction of the speech envelope

M100 amplitude. The amplitude of the 100 ms latency peak (M100, or N1m) was significantly higher in older adults than younger (F_[1,30]_ = 6.45, *p* = 0.017 ONH vs YNH and U(31) = 51, Z = - 2.699, *p* = 0.006 OHI vs YNH). No significant differences were seen between ONH and OHI (F_[1,27]_ = 0.249, *p* = 0.622). Because these results were consistent with previous studies (Soros et al., 2009, Alain et al., 2014), they were not further analyzed.

Reconstruction of the attended speech envelope. The ability to reconstruct the low-frequency speech envelope from cortical activity is a measure of the fidelity of the neural representation of that speech envelope (Ding et al., 2012, Presacco et al., 2016b). Figure 5 shows the correlation values for each single individual tested (top) along with the z-score of M100 (inset) and the grand average ± standard error of the reconstruction accuracy (bottom) for YNH, ONH and OHI for all the noise levels tested. All of the reconstruction values were significantly higher than the noise floor (all *p*-values < 0.01).

**Fig 5.**
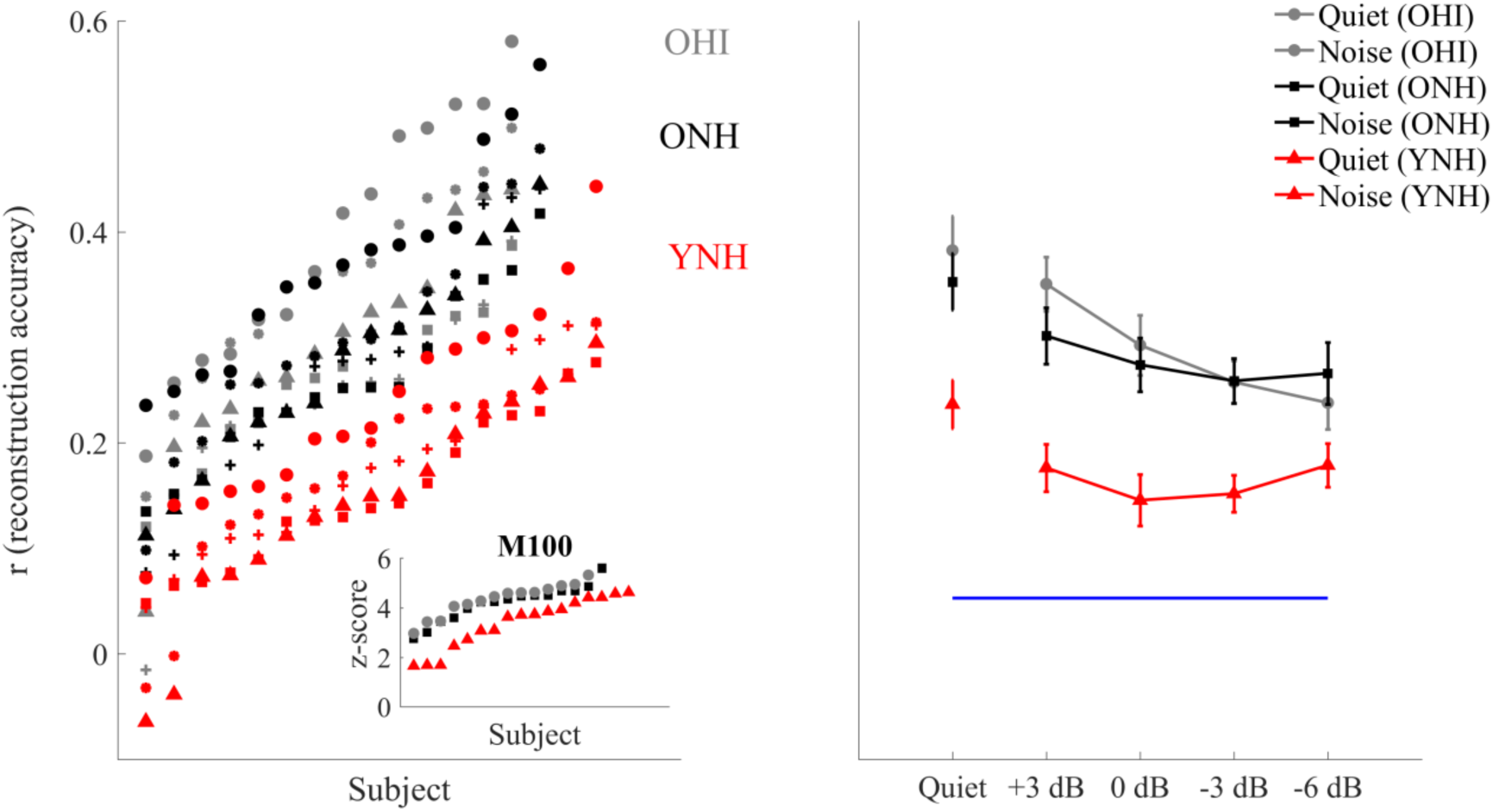
Left: plots of *r* values for each YNH (red), ONH (black) and OHI (gray) participant at each condition tested plotted in ascending order. The inset shows the z-score of the M100 recorded in response to a 500 Hz tone. Right: Reconstruction accuracy value 1 ± SE of the speech envelope of the foreground for YNH, ONH and OHI in quiet and in noise. The bottom horizontal line shows the noise floor. Older adults’ reconstruction fidelity is significantly better than that of the younger adults at all of the noise levels tested, but at -6 dB in OHI. Similarly, the M100 amplitude in both older adult listeners is significantly higher than younger adults. No significant differences and no interactions were found between ONH and OHI.

*Quiet:* One-way ANOVA showed significant differences among the three groups in quiet (F_[2,43]_ = 9.926*, p* < 0.001). Follow-up *t-*tests showed higher reconstruction values in ONH compared to YNH *(p* = 0.005) and in OHI compared to YNH and (*p* = 0.001) but not between ONH and OHI (*p* = 0.817).

*Noise:* Repeated measures ANOVA showed a main effect of group across SNRs (F_[2,43]_ = 11.567*, p* < 0.001). Follow-up one-way ANOVA showed higher reconstruction values in ONH compared to YNH (F_[1,31]_ = 16.563, *p* < 0.001) and in OHI compared to YNH (F_[1,28]_ = 21.815, *p* < 0.001), but not between ONH and OHI (F_[1,28]_ = 0.090, *p* = 0.767). There was a significant condition × group interaction (F_[2,43]_ = 3.149*, p* = 0.006) driven by a significant decline in reconstruction values with decreasing SNR in the OHI (*p* = 0.005) that was not seen in YNH (*p* = 0.525) or ONH (*p* = 0.141).

*Effect of the integration window*: The fidelity of the reconstruction was also tested at different integration windows, as described in Presacco et al., 2016b and the correspondent correlation values are shown in Figure 6. After correcting the *p*-values for multiple comparisons, results from Repeated Measures ANOVA showed significant differences between integration windows only in ONH (F_[2,28]_ = 14.954, *p* = 0.001, F_[2,28]_ = 20.457, *p* < 0.001, F_[2,28]_ = 4.897, *p* = 0.039, F_[2,28]_ = 10.489, *p* = 0.001, F_[2,28]_ = 5.048, *p* = 0.037 in quiet, +3, 0, -3 and -6 dB respectively), while no significant differences where found in YNH and OHI (all p-values > 0.05).

**Fig 6.**
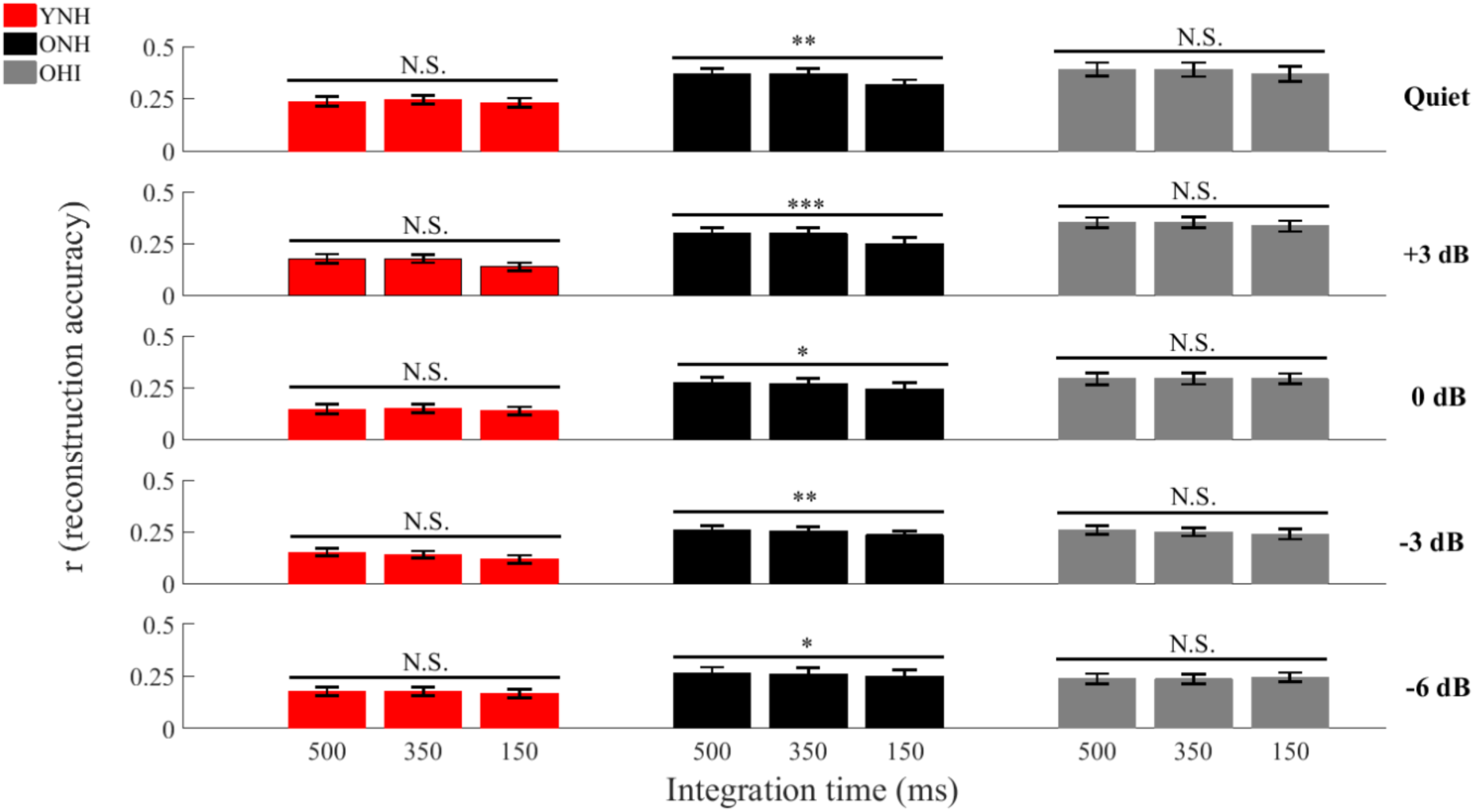
Reconstruction accuracy in quiet and at all the noise conditions tested for the 3 integration windows tested: 500, 350, and 150 ms. Significant differences across the 3 integration windows were found only in ONH in both quiet and at all the noise conditions tested. The size of the integration window seems to be playing a critical role only in older adults with normal hearing. \**P <* 0.05, \*\**P <* 0.01, \*\*\**P <* 0.001.

*Reconstruction of the unattended speech envelope.* One-way ANOVA showed no significant differences between ONH and OHI (all *p*-values > 0.05), while significantly higher correlation values were found in ONH with respect to YNH and OHI with respect to YNH in all the noise conditions tested but +3 dB (all p-values < 0.05).

### 3.6 Relationships among cognitive, midbrain and cortical data

The Flanker Inhibitory Control and Attention test showed significant differences among the three groups (F_[1,42]_ = 20.516, *p* < 0.001). Follow-up t-tests showed that YNH had significantly high scores than ONH (t_[30]_ = 5.232, *p* < 0.001) and OHI (t_[28]_ = 4.859, *p* < 0.001), while no significant differences were found between ONH and OHI (t_[26]_ = 0.05, *p* = 0.961). The Flanker score was evaluated with respect to the brain measures. Significant negative correlations (lower score associated with higher reconstruction accuracy) were found between the Flanker Inhibitory Control and Attention test score and the cortical response (average cortical decoding accuracy across all the noise levels) (ρ = -0.621, p = 0.013) in ONH, but not in YNH (ρ = 0.431, p = 0.084) or in OHI (ρ = 0.429, *p* = 0.144). No significant correlations were found among the Flanker Inhibitory Control and Attention test score and midbrain responses (average quiet-to-nose correlation across all the noise levels in the steady-state region and average stimulus-to-response correlation across all the noise levels) in any the three groups tested (all *p* > 0.05). Similarly, no significant correlations were found among midbrain and cortical responses in YNH and ONH (*p* > 0.05). However, significant positive correlations were found in OHI between cortex and average midbrain quiet-to-noise correlations across all the noise levels in the steady-state region (ρ = 0.719, *p* = 0.004), but not between cortex and average midbrain stimulus-to-response correlation across all the noise levels (ρ = 0.336, *p* = 0.240).

## 4. Discussion

The results of our study support almost all the hypotheses that we posited. Although hearing loss affects speech understanding in noise, we saw no significant differences in encoding of speech signals in noise between ONH and OHI in either the midbrain or the cortex, with the exception of the integration window analysis, where we unexpectedly found differences between the two older listener groups. Furthermore, two additional results that did not meet our expectations were the absence of a significant negative correlation between cognitive performance and cortical responses and the significant correlation between cortex and midbrain observed in OHI, suggesting that peripheral hearing loss might alter the relationship between the two areas of the auditory system that we have investigated.

### 4.1 Midbrain (EEG): Amplitude analysis

We found significant differences between YNH and the two groups of older listeners in quiet, but no significant differences between ONH and OHI. Our findings are consistent with the results from Ananthakrishnan et al. (2016), who recorded FFRs in quiet in response to the speech syllable /u/ in normal hearing and hearing impaired participants and reported that despite a higher degree of degradation of the envelope response in subjects with peripheral hearing loss, no significant difference between the two age groups were found.

This is not to say that peripheral hearing loss does not cause any noticeable change in the subcortical response. In a similar study conducted in older adults, Anderson et al. (2013) found a higher representation of the envelope in participants with hearing loss. Interestingly, these differences were exacerbated by the presence of background noise. The differing results between the current study and that of Anderson et al. (2013) might be explained by the dissimilarities between the two studies. First, the stimuli used in the studies differed not only in length, but in the composition of the spectral components. For the current study we used a 170 ms /da/, which had a relatively low amplitude noise burst during the first 10 ms and had a fundamental frequency of 100 Hz. Conversely, the stimulus used by Anderson et al. (2013) was very short (40 ms), had a fundamental frequency that linearly rose from 103 to 125 Hz, and had a shorter noise burst (5 ms) that was higher in amplitude than the one we used. The higher-amplitude noise burst of the 40-ms /da/ may activate a larger population of neurons, and because the OHI would have broader tuning bandwidths (Florentine et al., 1980), the differences between ONH and OHI might be more pronounced. The group differences in the Anderson et al. 2013 study were more apparent when the /da/ was presented in noise rather than in quiet. The noise stimulus in that study was pink noise, and again, the OHI participants may have higher response amplitudes to pink noise than would be expected to a single talker. The lack of significant differences between the older groups in our study might also be explained by differences in the audiogram with respect to Anderson et al. (2013). Our two older listener groups had significantly different hearing thresholds at all the frequencies tested, but at 125 Hz in the right ear, while in Anderson et al. (2013), these differences were reduced at low frequencies and more importantly their older normal hearing participants had better pure-tone average (PTA) thresholds. If we assume that peripheral hearing loss may somehow disrupt the balance between subcortical inhibitory and excitatory mechanisms (Caspary et al., 1995, Walton et al., 1998, Caspary et al., 2005, Caspary et al., 2006, Parthasarathy et al., 2010, Parthasarathy et al., 2011), it is possible that the lack of significant differences in response amplitude between ONH and OHI could be explained by poorer ONH hearing thresholds in the current study compared to those of Anderson et al. (2013)’s ONH group. In fact, the envelope response in the ONH group of the current study may be somewhat exaggerated due to PTA thresholds that are towards the limit of the normal hearing range.

Our results might also appear to contradict previous findings that have demonstrated exaggerated amplitudes associated with hearing loss in animal models (Kale et al., 2010, Henry et al., 2014). However, two important differences with respect to our experiment may limit the comparisons with these animal models. The first one is the etiology of hearing loss. Sensorineural hearing loss in chinchillas was induced by noise overexposure, while our subjects did not report any history of unusually high noise exposure. Second, the increase in response amplitude in the Henry et al. (2014) study was observed only in fibers whose characteristic frequency was between 1 and 2 kHz. The envelope of our FFR had a fundamental frequency of 100 Hz, a frequency at which Henry et al. (2014) did not find any increase in amplitude response.

### 4.2 Midbrain (EEG): Robustness of the Envelope to Noise

The quiet-to-noise correlation and the stimulus-to-response correlation analyses showed different results between ONH and OHI.

*Quiet-to-noise correlation analysis*

YNH showed a significantly more robust response to noise than OHI in both the transition and steady-state regions, but no significant differences were found between YNH and ONH, or between ONH and OHI. Interestingly, in this analysis we saw significant declines in the robustness of the response across noise conditions in the transition and steady-state regions in YNH and ONH, but not in OHI, suggesting that peripheral hearing loss further deteriorates the latency of the encoded speech envelope already affected by aging (Pichora-Fuller et al., 2007, Anderson et al., 2012, Presacco et al., 2015, Mamo et al., 2016, Presacco et al., 2016a, b) to the point that even in relatively favorable conditions (e.g. +3 dB), the timing of the response is already significantly compromised. A prolongation in timing will affect the strength of the correlation between responses obtained in quiet and noise.

These results might be informed by those of Mehraei et al. (2016) who recorded auditory brainstem responses (ABRs) to click stimuli presented in quiet and in several levels of broadband noise. In young normal-hearing adults, the ABR Wave V latency is expected to increase with decreasing SNRs due to neural desynchronization. The slope of this latency increase was shallower, however, in some of the study participants who had lower Wave I amplitudes and poorer performance on a temporal processing task. Mehraei et al. surmised that the ABR latency in noise may be a marker of cochlear synaptopathy. Our findings showing a decrease in correlation values with increased noise in the YNH and ONH but not in the OHI participants might provide support for the idea that the age group differences for other FFR measures (amplitude and stimulus-to-response) are driven by central rather than peripheral factors. If the ONH group was significantly affected by peripheral hearing loss, synaptopathy, or loss of auditory nerve fibers, we might have expected to find a group × noise level interaction between ONH and YNH, but both groups experienced a similar loss of correlation value with increased noise levels.

*Stimulus-to-response correlation*

The YNH group had significantly higher stimulus-to-response correlations than either ONH or OHI, but there were no differences between ONH and OHI. These results contrast with the quiet-to-noise correlations that showed an overall effect of hearing loss but not of aging. The differences between the analyses may explain these seemingly conflicting results. The quiet-to-noise correlation is affected by the timing of the response, which is compromised by reduced audibility. The stimulus-to-response correlation is affected by the degradation of the neural speech envelope, and the analysis is independent of the latency delay. The fact that YNH had a better representation of the stimulus with respect to ONH and OHI and that no differences were found between the two older adults groups speaks in favor of an age-related degradation of the response, rather than a degradation associated with hearing loss.

### 4.3 Cortex (MEG): Reconstruction of the speech envelope

The results from the cortical analysis confirm the existence of overrepresentation of the response envelope in older adults with and without peripheral hearing loss. The reconstruction accuracy of OHI is significantly higher than YNH in both quiet and noise, a finding that is consistent with other studies showing an exaggerated cortical response associated with age (Tremblay et al., 2003, Soros et al., 2009, Lister et al., 2011, Alain et al., 2014, Presacco et al., 2016a, b). In our previous experiments (Presacco et al., 2016a, b), we speculated that this abnormally high cortical response could be due to a mix between an age-related imbalance between inhibitory and excitatory mechanisms and age-related cognitive deficits. However, we could not rule out the possibility that loss of hearing sensitivity could be the driving factor. Our current study suggests that indeed age may be the driving factor behind changes at the cortical level. We could not find any significant difference between ONH and OHI in any of the conditions tested, even including responses to the presentation of a simple tone. Had peripheral hearing loss been the driving factor in explaining this overrepresentation, we would have expected significantly higher reconstruction accuracy in OHI with respect to ONH. These findings were also confirmed in the analysis of the unattended speech, which was significantly higher in the two older listener groups with respect to the younger participants. The only surprising difference between ONH and OHI arose from the integration windows analysis. In our previous study (Presacco et al., 2016b), it was shown that narrowing the integration window negatively affected the reconstruction accuracy of ONH, but had no significant effect on YNH. We speculated that this result was consistent with previous psychoacoustic (Fitzgibbonsand Gordon-Salant 2001; Gordon-Salant et al. 2006) and electrophysiological (Alain et al. 2012; Lister et al. 2011) experiments showing that older adults had more problems adapting to changes in temporal parameters, and therefore expected to see similar results in OHI. The difference may arise due to mechanisms having been altered by hearing loss; several studies have shown that loss of auditory sensitivity may affect the reorganization of different areas of the brain, thus leading to different speech encoding strategies in ONH and OHI (Peelle et al., 2011, Campbell et al., 2013, Du et al., 2016, Vaden et al., 2016). More studies will be required to elucidate this unexpected result.

### 4.4 Relationships among cognitive function, midbrain and cortical responses

One of the most important and intriguing results of our previous experiment (Presacco et al., 2016a) was the lack of correlation between midbrain and cortex and a significantly negative correlation between cortex and cognitive scores in ONH, in contrast to the significant correlation between cortical and midbrain responses, and lack of correlation between cortex and cognitive scores, seen here for OHI. We speculated that the lack of correlation was consistent with results from a recent animal study (Chambers et al., 2016) that showed the existence of compensatory central gain increases that may restore the representation of the auditory object in cortex even when the input from the brainstem is severely degraded. These results underscored the importance of studying auditory temporal processing at different levels of the auditory system to better understand how degraded encoding of the signal at lower levels of the auditory system may affect the final representation of speech in the cortex. The negative correlation between cortex and cognitive scores supported our hypothesis that an overrepresentation in the cortex was not a biomarker that represents an advantageous response of the brain but rather, an abnormally high neural activity that could indicate failure in processing auditory information, as also reported by a recent study (Millman et al., 2017). The presence of peripheral hearing loss was therefore a critical factor for this analysis, because several studies have shown how loss of auditory sensitivity affects the reorganization of different areas of the brain (Peelle et al., 2011, Campbell et al., 2013, Du et al., 2016, Vaden et al., 2016). Our results indeed show a different relationship between midbrain and cortex in OHI, as indicated by a significant positive correlation between cortical and midbrain responses. This finding may suggest that peripheral hearing loss is associated with higher interdependence between midbrain and cortex, consistent with the results from Bidelman et al. (2014). In Bidelman’s study this correlation was seen only in the older subjects, many of whom had significant hearing loss, and it was not seen in the younger subjects.

Although the ONH group showed a negative correlation between the Flanker Inhibitory Control and Attention test score and the reconstruction value of the cortical envelope, neither the YNH nor the OHI groups showed a signification relationship between the two measures; in fact, the direction of the correlation for both YNH and OHI groups was in a positive rather than in a negative direction. This was an unexpected finding for the OHI group given our theory that overrepresentation of the cortical response does not indicate enhanced temporal processing. We argue that a feasible explanation exists to reconcile these incongruent results. We previously noted a midbrain-cortical association in the OHI that was not seen in the ONH, suggesting a tighter link between subcortical and cortical regions in individuals with hearing loss (Bidelman, 2014). Similarly, in cortex, hearing loss leads to reorganization, with a loss of gray matter volume in primary auditory cortex (Peelle et al., 2011) and higher activation of frontal cortical areas (Campbell et al., 2013, Du et al., 2016, Peelle et al., 2016). Therefore, it is possible that hearing loss affects cognitive-cortical relationships differently than the presence of aging alone.

## 5. Conclusion

The overall results of our study bring additional support to the hypothesis that central temporal auditory deficits are critical factors in the communications problems experienced by older adults. Subcortical and cortical responses alone show no significant differences between the two older listener groups, suggesting that aging is a driving factor in explaining degradation of speech comprehension in noise. What remains to be elucidated from this experiment are the significant correlation between cortical and midbrain responses in the participants with hearing loss and the lack of sensitivity to changes in the integration window in OHI. At this point, we can only speculate that this relationship is due to functional changes in the brain caused by the presence of hearing loss. Future directions will focus on using the biomarkers identified in this study to assess the efficacy of auditory training techniques. Specifically, it will be critical to understand if an improvement in speech understanding is associated with changes in the neural response, such as reduced degradative effects of noise on midbrain responses and reduced overrepresentation of the cortical response.

## ACKNOWLEDGEMENT

Funding for this study was provided by the University of Maryland College Park (UMCP) Department of Hearing and Speech Sciences, UMCP ADVANCE Program for Inclusive Excellence (NSF HRD1008117), and National Institute on Deafness and Other Communication Disorders (R01DC014085, T32DC-00046 and T32DC-010775-07).

We would like to thank Katlyn B. Van Dyke for her help in collecting data and Natalia Lapinskaya for excellent technical support.

Hear Res.

